# Detection of domain motion in NADPH-cytochrome P450 oxidoreductase through polarization anisotropy measurements

**DOI:** 10.1101/178426

**Authors:** Elizaveta A Kovrigina, Chuanwu Xia, Jung-Ja P. Kim, Evgenii L Kovrigin

**Affiliations:** Biochemistry Department, Medical College of Wisconsin, Milwaukee, WI 53226; Chemistry Department, Marquette University, Milwaukee, WI 53201

## Abstract

Conformational transitions between closed and open states in the NADPH-cytochrome P450 oxidoreductase (POR) play a critical role in its electron-transport function. In this study, we determined rotational diffusion coefficients of the EDANS fluorophore attached to the cytosolic POR construct lacking the N-terminal transmembrane region. We identified two dynamic modes, slow and fast, which are interpreted as the rotational diffusion of POR as a whole and the local domain motion, respectively. Timescale of the local rotational diffusion component suggests that it may correspond to the transient opening of the fully oxidized POR structure.

---

NADPH-cytochrome P450 oxidoreductase (POR) transfers electrons from NADPH to multiple cytochromes P450 and other proteins in the endoplasmic reticulum of liver cells and other tissues^*1*^. Conformational changes in POR molecule are required for productive interaction with cytochromes P450, and the "open" and "closed" structures of POR were identified by X-ray crystallography^*2-4*^. Because the complex of POR with cytochrome P450 has never been structurally resolved, how these conformational states of POR relate to the steps in the functional cycle—remains debated. For example, oxidized POR has been demonstrated to assume closed conformation (blue ribbon in Figure 1) in solution by NMR spectroscopy and small-angle X-ray scattering^*5-7*^, while other reports indicated it is in an open form in the oxidized state^*8, 9*^. In this study, we probed conformational states of the oxidized cytosolic fragment of POR, residues 57-678, (in the following text—POR) from viewpoint of the local conformational freedom measured by rotational diffusion constants. We used a single cysteine mutant of POR with otherwise cysteine-less background to introduce a long-lived EDANS fluorophore in the FMN domain of POR. Two distinct time constants observed in fluorescence anisotropy decays allowed us to propose that FMN domain might have some degree of local mobility within the oxidized POR structure.

**Figure 1.**
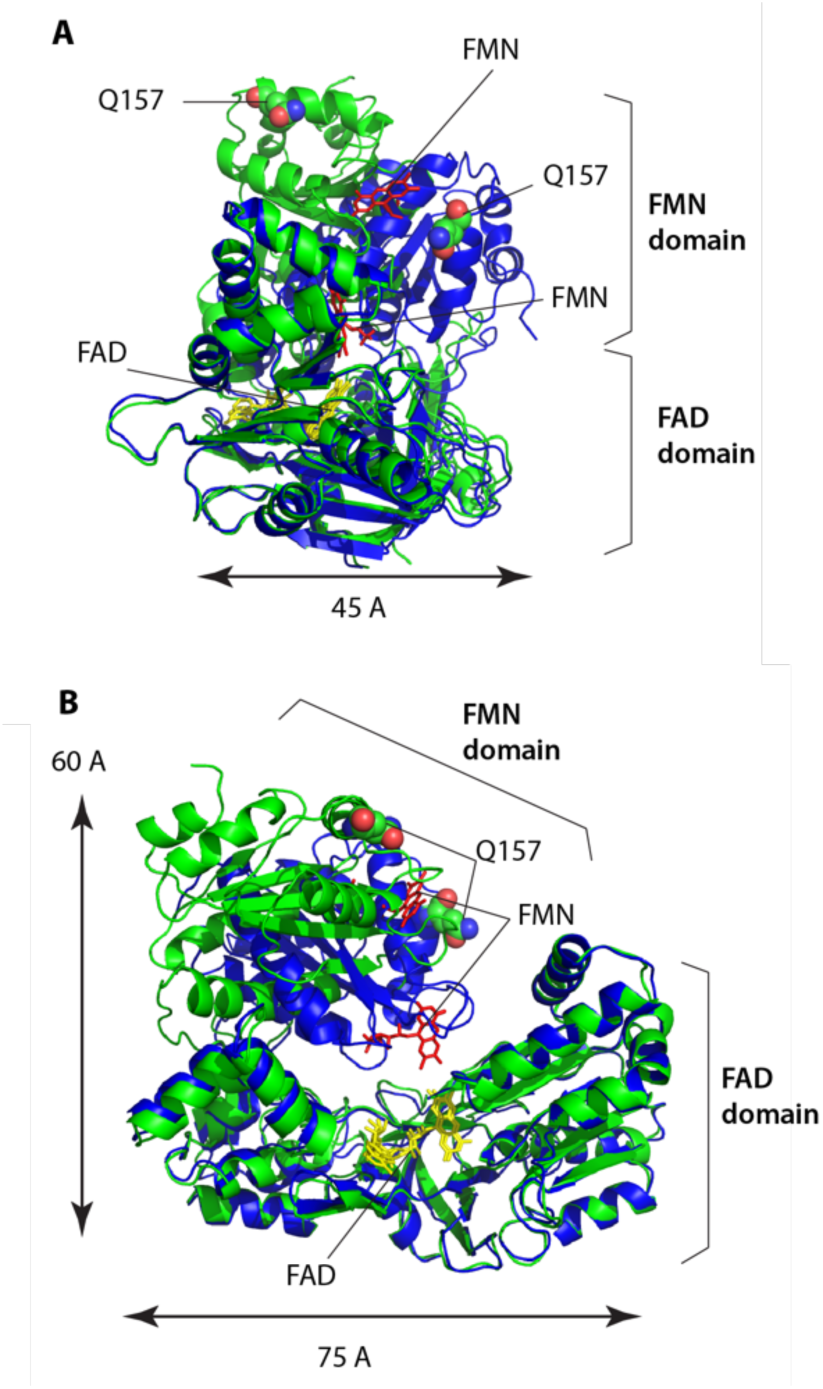
Cytosolic fragment of POR, residues 57-678: the closed conformation (blue ribbon, PDB ID 1AMO) and the putative open conformation (green ribbon, PDB ID 3ES9). Panel **B** is 90-degree rotation of **A** around the vertical axis. The structures have the FAD domain aligned. FMN cofactor, red; FAD cofactor, yellow. Q157, spheres.

## Results and Discussion

Figure 1 shows an alignment of the closed oxidized POR^*2*^ (represented with blue ribbon) and the hypothetical open state modeled by a conformationally restrained mutant with the four-residue deletion in the interdomain loop (TGEE at the positions 236-239; in the following—ΔTGEE)^*4*^. The two structures are aligned by the larger domain—FAD domain, to highlight a possible motion of the FMN domain that represents the opening transition necessary for interaction with the cytochrome P450.

Viewing this protein structure from the mechanistic standpoint, closed state (blue) represents a well bound conformation that is largely stabilized by interdomain contacts. The conformation of ΔTGEE mutant (green) shows these contacts broken and the FMN domain orientation, likely, stabilized by crystal contacts. Therefore, as soon as the closed conformation dissociates the non-covalent bonds at the interdomain interface, one might expect relatively free local rotational diffusion of the FMN domain with respect to the rest of the POR molecule. Therefore, measurements of local rotational correlation times might be informative on the conformational equilibrium in POR.

Fluorescence polarization anisotropy measurement allows for a sensitive detection of rotational diffusion of fluorophores attached to proteins^*10*^. To allow for accurate measurements, the life time of the fluorescent probe should be long enough—comparable to the rotational correlation time of the protein. In this study, we utilized [5-((2-aminoethyl)amino) naphthalene-1-sulfonic acid (EDANS) fluorophore due to its appropriately long life time (14 ns). Site-specific attachment of the fluorophore to the surface of POR was achieved by using the POR construct with all native cysteines mutated out (cysless POR)^*11*^ and a single surface-exposed cysteine introduced at the position 157 in the FMN domain. EDANS-maleimide conjugate was attached to C157 via an irreversible reaction of maleimide and thiol groups.

To remove protein aggregates as well as excess of unreacted EDANS from the POR samples before fluorescence measurements, the labeling reaction mixtures were separated using size-exclusion chromatography. The EDANS-labeled POR eluted as a single peak in the expected molecular weight range of 70 kDa.

We noted that any attempts to concentrate labeled POR resulted in aggregation that manifested itself as an extremely long-lived polarization component. Therefore, to exclude formation of aggregates, the samples from size-exclusion profile were used for polarization anisotropy measurements directly. Typical protein concentrations were in the range of 2-5 μM, and the fluorescence decays were collected for 40-50 hours to accumulate sufficient signal.

To verify specificity of labeling we also used the "parent" cysless POR construct in the labeling reaction with EDANS maleimide at the same conditions as for the Q157C mutant. Purified cysless POR showed specific EDANS emission of 1.2 x10^5^ counts sec^−1^ (μM POR)^−1^, while the Q157C POR produced 14 x10^5^ counts/sec^−1^ (μM POR)^−1^. Therefore, the non-specifically bound EDANS contributed less than 10% of the fluorescence signal in the Q157C-EDANS POR samples.

Figure 2 shows the polarization anisotropy decay of the Q157C-EDANS POR. Fitting with Eq. 1 revealed that the EDANS fluorescence anisotropy decays with two major modes characterized by the short and long rotational correlation times, t1 and t2, respectively. Large uncertainty of the measured correlation times does not allow quantitative modeling of the motions in POR but may be used to attribute the motions to the global tumbling or local mobility.

**Figure 2.**
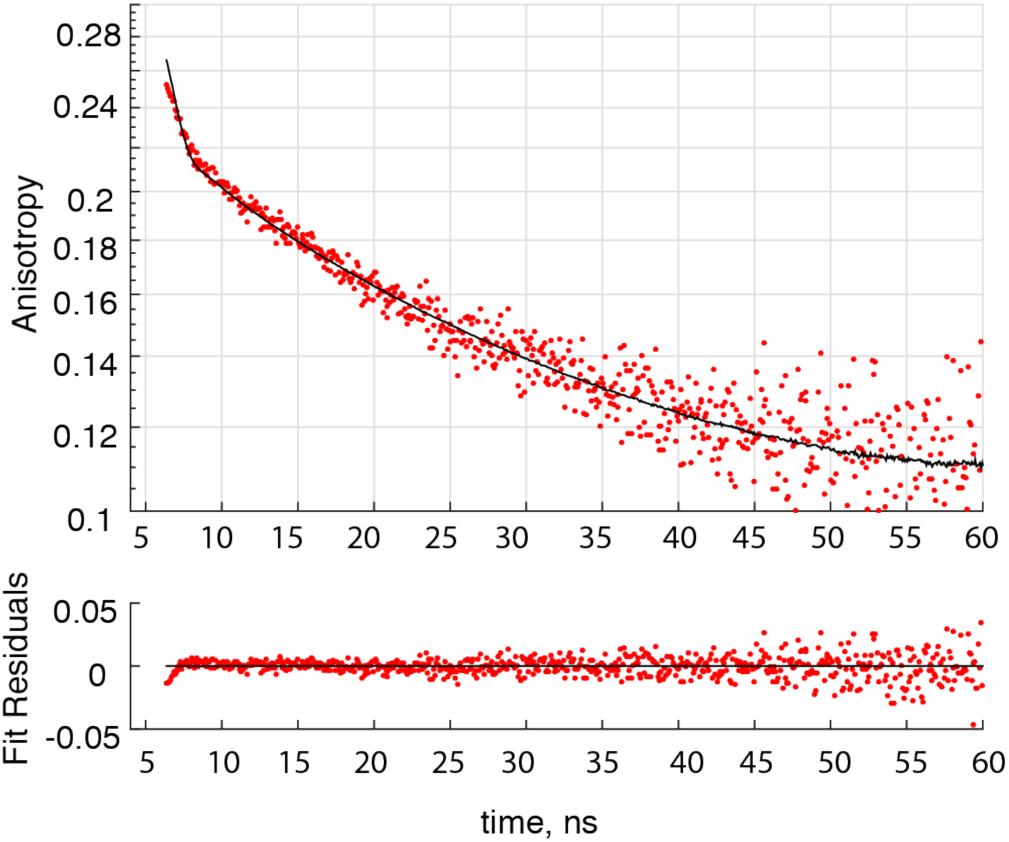
Fluorescence polarization anisotropy decay (top) and residuals from fitting (bottom) for Q157C POR labeled with EDANS-maleimide. Experimental data, red dots; best-fit curve, black solid line. Fitter parameters of Eq. 1 with uncertainty estimates as reported by *AniFit*: r_0_ = 0.18 [ 0.14 - 0.22 ], b_1_ = 0.03 [ < 0.1 ], t_1_ = 9 [ < 34] ns, t_2_ = 50 [ 10 - 90 ] ns.

The molecular shape of the cytosolic POR construct shown in Figure 1 may be roughly approximated as an oblate ellipsoid with the axis ratio 5/7. Considering its molecular weight of 71 kDa, we calculated overall rotational correlations times in excess of 33 ns. This estimate allows us to speculate that the longer life time component, t2, originates from the overall rotational diffusion of the POR molecule as a whole. The faster component, t1, is unlikely to originate from depolarization of the EDANS fluorophore due to its own reorientation on the linker because rotational diffusion correlation times of small molecule (segmental motion) is, typically, very short—in the sub-nanosecond range. In Q157C POR, the t1 is around 9 ns, which likely represents the *local dynamics* of the POR structure (for a comparison, 9 ns would be a correlation time of a 20 kDa spherical protein). Whether this local rotational freedom corresponds to FMN domain undocking from FAD domain as shown in Figure 1 remains to be seen. To establish this, our future work will include mutants introducing EDANS label in the FAD domain and use of the "open" ΔTGEE POR as a reference.

## Methods

The cytosolic cysless POR constructs were prepared as described in our earlier report^*12*^. Size-exclusion chromatography was carried out using Superose 6 Increase 10/300 column on the Shimadzu HPLC system. Polarization anisotropy decays were recorded with QM40 QuantaMaster (HORIBA Scientific, Edison, NJ) equipped with PicoMaster 1 TCSPC following the published protocol^*13*^ at 20°C. Fluorescence data were analyzed with *AniFit* software (kindly shared by Søren Preus; available from www.fluortools.com). Polarization anisotropy was fit with a three-exponential function:

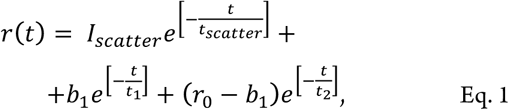

where the first term accounts for the scattered light contribution while the polarization anisotropy decay is described by the two additional exponential terms^*10*^. The protocol for simulations of correlation times of the POR molecule followed Kawski^*14*^ and Lakowicz^*10*^ and was described in detail previously^*13*^. The MATLAB script to perform calculations of rotational correlation time for ellipsoids is included in Supporting Information.

## Acknowledgement

This work was supported in part by NIH GM097031 to JJK.

## References

[1] Waskell, L., and Kim, J.-J. P. (2015) Electron Transfer Partners of Cytochrome P450, In Cytochrome P450. Structure, Mechanism, and Biochemistry (Ortiz de Montellano, P. R., Ed.) 4th ed., pp 33–68, Springer Cham Heidelberg New York Dordrecht London.

[2] Wang, M., Roberts, D. L., Paschke, R., Shea, T. M., Masters, B. S., and Kim, J. J. (1997) Three-dimensional structure of NADPH-cytochrome P450 reductase: prototype for FMN- and FAD-containing enzymes, Proc Natl Acad Sci U S A 94, 8411–8416.

[3] Sugishima, M., Sato, H., Higashimoto, Y., Harada, J., Wada, K., Fukuyama, K., and Noguchi, M. (2014) Structural basis for the electron transfer from an open form of NADPH-cytochrome P450 oxidoreductase to heme oxygenase, Proceedings of the National Academy of Sciences 111, 2524–2529.

[4] Hamdane, D., Xia, C., Im, S. C., Zhang, H., Kim, J. J., and Waskell, L. (2009) Structure and function of an NADPH-cytochrome P450 oxidoreductase in an open conformation capable of reducing cytochrome P450, J Biol Chem 284, 11374–11384.

[5] Ellis, J., Gutierrez, A., Barsukov, I. L., Huang, W.-C., Grossmann, J. G., and Roberts, G. C. K. (2009) Domain Motion in Cytochrome P450 Reductase. Conformational Equilibria Revealed by NMR and Small-Angle X-Ray Scattering, Journal of Biological Chemistry 284, 36628–36637.

[6] Vincent, B., Morellet, N., Fatemi, F., Aigrain, L., Truan, G., Guittet, E., and Lescop, E. (2012) The closed and compact domain organization of the 70-kDa human cytochrome P450 reductase in its oxidized state as revealed by NMR, J Mol Biol 420, 296–309.

[7] Frances, O., Fatemi, F., Pompon, D., Guittet, E., Sizun, C., Perez, J., Lescop, E., and Truan, G. (2015) A well-balanced preexisting equilibrium governs electron flux efficiency of a multidomain diflavin reductase, Biophys J 108, 1527–1536.

[8] Hedison, T. M., Hay, S., and Scrutton, N. S. (2015) Realtime analysis of conformational control in electron transfer reactions of human cytochrome P450 reductase with cytochrome c, FEBS J 282, 4357–4375.

[9] Hedison, T. M., Hay, S., and Scrutton, N. S. (2017) A perspective on conformational control of electron transfer in nitric oxide synthases, Nitric Oxide-Biol. Chem. 63, 61–67.

[10] Lakowicz, J. R. (2010) Principles of Fluorescence Spectroscopy, 3rd ed., Springer.

[11] Xia, C., Hamdane, D., Shen, A. L., Choi, V., Kasper, C. B., Pearl, N. M., Zhang, H., Im, S. C., Waskell, L., and Kim, J. J. (2011) Conformational changes of NADPH-cytochrome P450 oxidoreductase are essential for catalysis and cofactor binding, J Biol Chem 286, 16246–16260.

[12] Kovrigina, E. A., Pattengale, B., Xia, C., Galiakhmetov, A. R., Huang, J., Kim, J.-J. P., and Kovrigin, E. L. (2016) Conformational states of cytochrome P450 oxidoreductase evaluated by FRET using ultrafast transient absorption spectroscopy, Biochemistry 55, 5973–5976.

[13] Kovrigina, E. A., Galiakhmetov, A. R., and Kovrigin, E. L. (2015) Ras G domain lacks intrinsic propensity to form dimers, Biophysical Journal 109, 1000–1008.

[14] Kawski, A. (1993) Fluorescence anisotropy: theory and applications of rotational depolarization, Crit Rev Anal Chem 23, 459–529.

